# Fluorescence Shadow Imaging of *Hypsibius exemplaris* Reveals Morphological Differences Between Sucrose- and CaCl_2_-Induced Osmobiotes

**DOI:** 10.1101/2024.02.14.580268

**Authors:** Brendin B. Flinn, Hayden M. O’Dell, Kara M. Joseph, Amanda L. Smythers, David P. Neff, Leslie M. Hicks, Michael L. Norton, Derrick R.J. Kolling

**Affiliations:** Marshall University, Department of Chemistry, Huntington, WV, USA; University of North Carolina at Chapel Hill, Department of Chemistry, Chapel Hill, NC, USA

**Keywords:** Tardigrades, Cryptobiosis, Volume, Microscopy

## Abstract

Tardigrades are renowned for their ability to survive a wide array of environmental stressors. In particular, tardigrades can curl in on themselves while losing a significant proportion of their internal water content to form a structure referred to as a tun. In surviving varying conditions, tardigrades undergo distinct morphological transformations that could indicate different underlying mechanisms of stress sensing and tolerance specific to the stress condition. Presently, methods to effectively distinguish between morphological transformations, including between tuns induced by different stress conditions, are lacking. Herein, a new approach for discriminating between tardigrade morphological states is developed and utilized to compare sucrose- and CaCl_2_-induced tuns. A novel method of shadow imaging with confocal laser scanning microscopy enabled production of three-dimensional renderings of tardigrades in various physiological states resulting in tardigrade volume measurements. Combining these measurements with qualitative morphological analysis using scanning electron microscopy revealed that sucrose- and CaCl_2_-induced tuns have distinct morphologies, including differences in the amount of water expelled during tun formation. Further, varying the concentration of the applied stressor did not affect the amount of water lost, pointing towards water expulsion being a controlled process that is adapted to the specific stressors.

**Summary Statement:** A novel method for volume measurements of biological specimens was developed and paired with scanning electron microscopy to allow for morphological comparisons between tardigrade osmobiotes.

## 1. Introduction

Tardigrades are microscopic animals that comprise the minor phylum Tardigrada.^1^ Interest in tardigrades is motivated by their unique extremotolerant characteristics that allow them to survive a wide array of environmental stressors, many of which could be leveraged for therapeutic use by humans.^2^ A non-exhaustive list of conditions tardigrades can survive includes extreme temperatures (as high as over 100 °C and as low as -272 °C, just one degree above absolute zero), extreme pressures (as low as the vacuum pressure of space and as high as 600 MPa), prolonged starvation, desiccation, and intense radiation.^3-9^ These environmental stressors can induce cryptobioses, states characterized by significantly downregulated metabolic activity. Tardigrades respond to many of these stressors by curling their body in on themselves while decreasing their internal water content to form a compact structure referred to as a tun, a process through which tardigrades can survive extreme conditions for prolonged periods of time.^10^

Prior work has thoroughly discriminated broad morphological differences between cryptobioses, such as describing the phenomena through which desiccated tardigrades form a tun while tardigrades in generally deteriorating environments form cysts.^10^ However, work to date has not rigorously defined fine morphological differences between specimens exposed to similar conditions, such as discriminating between tuns induced by diverse stressors. Beyond a limited focus on comparing differences in morphology, the only reported means of quantifying a morphological feature other than linear dimensions by microscopy is an approximation of tardigrade volume that estimates tardigrades as either cylinders (in their hydrated state) or hemicylinders (in tun states).^11^ This approach is not suitable for comparative analysis between states because tardigrades are not geometrically regular shapes (e.g., they are not smooth-surfaced, their anterior and posterior regions taper and do not approximate circles, and they have limbs protruding from their main body column). Thus, using the same approximation for states that have apparent morphological differences would introduce measurement error into one group more than it would to the other.^11^ An improved approach was therefore implemented in which the leg volume is calculated separately from the main body volume; however, this still propagates error due to deviations in tardigrade morphology from cylindrical features.^12^ Another approach implemented X-ray computed tomography which had the benefit of not employing geometric approximation but requires a custom-built instrument to image a sample the size of a tardigrade while maintaining high resolution. Additionally, this approach was only utilized for a single specimen, thus its quantitative figures of merit and scalability are unknown.^13^ Thus, there are currently limited methods by which distinct morphological changes can be identified and/or quantified.

It has long been observed that tun formation requires active processes to protect against damage in desiccating environments, and more recent investigations have extended these observations to osmotic stressors as well.^11, 14-18^ For instance, acclimation periods to low concentrations of osmolytes can allow for increased survival upon transfer to higher concentrations of the same osmolytes, indicating that tardigrades require time to carry out physiological processes of protection to enable survival.^19^ We and others have observed that tuns induced by exposure to high concentrations of ionic and non-ionic osmolytes have apparent distinct morphological features as well as differences in survivability.^18, 19^ Given the active nature of tun formation and that tardigrades exist in a plethora of different environments, differences in tun morphology in response to different osmotic stressors may be indicative of distinct mechanisms of stress sensing acquired as adaptations to varying environments. However, approaches to discriminate between tuns induced by individual stress conditions are lacking. Thus, in order to discern distinct morphological changes between stress conditions, it is necessary to develop an accessible, scalable, and quantitative method for rigorous morphological characterization. These morphological characterization methods may aid in elucidating differential responses to different stressors, providing insight into the different ways that tardigrades may sense and respond to environmental changes.

To achieve detailed morphological characterization, confocal laser scanning microscopy (CLSM) and scanning electron microscopy (SEM) were employed in combination. A novel use of fluorescence-based shadow imaging was developed that generated three-dimensional renderings from which body volumes could be calculated, enabling quantification of volume differences between different tardigrade states. SEM allowed for qualitative observations of structural differences between cryptobiotes. Combining the observation of morphological features through SEM and the measurement of total body volumes through shadow imaging, it was possible to discriminate between sucrose- and CaCl_2_-induced tuns, demonstrating both qualitatively and quantitatively that the two states are distinct from one another. This demonstrates that *Hypsibius exemplaris* is responding to each stressor differently despite both being putative osmotic stressors.

## 2. Materials and Methods

### 2.1 Tardigrade Husbandry

All-female cultures of parthenogenically reproducing *H. exemplaris* (Sciento, Manchester, UK) were reared in 1-or 2-L Erlenmeyer flasks on stationary phase *Chlorella vulgaris* (unicellular algae). Culture medium was changed biweekly using 40-µm Corning™ Sterile Cell Strainers (Sigma-Aldrich, St. Louis, MO), which retains the tardigrades while allowing algae and waste to flow through. The filter was inverted and rinsed into the same flask with the addition of fresh deionized water. Tardigrades were fed weekly with *C. vulgaris* that was grown photoautotrophically under previously established protocols.^20^ Cultures were maintained at room temperature on a 12:12 h light:dark cycle using a 7 W (630 lumens) LED lamp.^20^

### 2.2. Cryptobiosis Induction and Anesthetization

To induce sucrose and CaCl_2_ tuns, tardigrades were collected in glass petri dishes with minimal algae present and swirled to the center of the dishes to form large masses of tardigrades. Approximately 10 µL of tardigrades were collected for each trial and transferred to new, empty glass petri dishes. The tardigrades were then immediately dosed with working concentrations of 600 mM sucrose (Sigma-Aldrich, St. Louis, MO) for 1 h, 75 mM CaCl_2_ (AMRESCO, Solon, OH) for 5 h, or 12.5% (v/v) methanol (Thermo Fisher Scientific, Waltham, MA). Methanol dosing was used to prevent movement of hydrated specimens during imaging. Optimal concentrations of stressors and exposure times were determined in prior work.^18^ The concentration of sucrose was varied (300 mM, 450 mM, and 600 mM) in a separate experiment to test for concentration-dependent effects on water expulsion during tun formation; tardigrades were exposed to each concentration for 3 h, 1 h, and 1 h, respectively.^18^

### 2.3. Scanning Electron Microscopy

Tardigrades in each of the three states mentioned in *4.2* were collected in the center of their glass petri dishes and pipetted with minimal volume into new glass petri dishes containing 4% (w/v) formaldehyde (Thermo Fisher Scientific, Waltham, MA) for fixation. All samples were fixed overnight at 4 °C. After fixation, the tardigrades were pipetted in minimal volume into new glass petri dishes containing deionized water; this was repeated two more times. After rinsing off the formaldehyde solution, all samples were dehydrated with an increasing ethanol (Thermo Fisher Scientific, Waltham, MA) series (50, 70, 80, 90, and 3x 100% (v/v) ethanol) with a 5-min incubation in each concentration of ethanol. After the final rinse in 100% (v/v) ethanol, 10 µL of each sample of fixed and dehydrated tardigrades was pipetted onto separate aluminum stubs (into the direct center in order to avoid any tardigrades being washed off of the sides of the stubs). The stubs were then immediately placed in a Polaron SEM coating system (Quorum Technologies, Lewes, UK) and were coated with Au-Pd for 3 min. All sample stubs mounted with tardigrades were imaged on a JEOL JSM 7200FLV (JEOL, Tokyo, Japan) scanning electron microscope with a spot size of 10 and working distance of 10 mm. Image magnification varied but is labelled on all SEM micrographs presented.

### 2.4. Shadow Imaging (Confocal Laser Scanning Microscopy)

This protocol was adapted from the SUSHI (super resolution shadow imaging) technique recently described for the measurement of extracellular fraction volume in unlabeled brain tissue.^21^

Tardigrade samples were prepared as described in *2.2*. Ten microliters of each sample (containing 10-20 tardigrades) was pipetted onto a glass slide (Thermo Fisher Scientific, Waltham, MA) outfitted with a 120 µm deep spacer well (Invitrogen, Waltham, MA). One microliter of 500 µM cell-impermeant calcein (ex./em. maxima of 494/514 nm) (Thermo Fisher Scientific, Waltham, MA) stock solution in dimethyl sulfoxide (DMSO) (Thermo Fisher Scientific, Waltham, MA) was pipetted into the center of the sample droplet and mixed in with the pipette tip. The spacer well was covered with a glass no. 2 coverslip (Thermo Fisher Scientific, Waltham, MA) and immediately placed on the specimen stage of the confocal microscope. All images were captured on a Leica SP5 TCSII (Leica, Wetzlar, Germany) outfitted with an argon laser as well as a Coherent Chameleon Vision II multiphoton laser (Coherent, Santa Clara, CA) (tunable IR laser emitting between 680 and 1080 nm).

Thirteen tardigrades were imaged for each condition using either the 496 nm line of the argon laser or the multiphoton (MP) laser tuned to 920 nm (with 50% transmission) as an excitation source. When using the argon laser for excitation, an adjustable photomultiplier tube set to detect between 510-520 nm was used for fluorescence detection. When using the MP laser for excitation, a 500-550 nm band pass non-descanned detector (NDD PMT) was used. The following parameters were used for image acquisition: objective, 20x dry; zoom factor, 1.5; resolution, 1024×1024 pixels; z-stack step distance, 0.5 µm. Brightfield images were collected in tandem for all specimens. Samples were excluded from analysis if the renderings produced via shadow imaging did not represent the whole specimen or otherwise did not accurately depict the specimen (e.g., missing limbs or incomplete rendering due to proximal air pockets). Thirteen specimens were imaged for each group to ensure that a minimal sample size of 10 was acquired after any necessary exclusions.

All image processing/analysis was conducted in the FIJI port of ImageJ.^22^ Image files were exported into ImageJ and the images were inverted and converted to binary (to set the pixels comprising the tardigrade at an intensity of 255). From the binarized images, 3D renderings of the tardigrades were produced using the 3D viewer plugin and compared to the images from the transmitted light detector to ensure accurate renderings were acquired. The 3D objects counter plugin was used to compute the volume of each tardigrade.^23^ Statistical significance between conditions was determined using Welch’s t-test. Brightfield images of specimens from each condition were used for measurements of the following linear body parameters: anterior to posterior length, lateral width, and width from dorsal to ventral sides. Measurements were conducted in the FIJI port of ImageJ by drawing lines along the specified regions and measuring the length of those lines.

#### 2.4.1. Single-Specimen Shadow Imaging

To test if averaged volume changes from population measurements were comparable to the volume changes of individual organisms, samples of single sucrose-dosed tardigrades were prepared for imaging as described in *2.4* and then allowed to rehydrate in distilled water. The rehydrated specimens were then prepared for imaging as described for hydrated specimens in *2.4*. The % change in volume between the hydrated and sucrose-induced tun state was calculated for each specimen individually and averaged before being compared to the % difference between the average volumes of the hydrated and sucrose-dosed specimens.

## 3. Results

### 3.1. Scanning Electron Microscopy Reveals Morphological Differences Between Osmobiotes

No consistent appearance of gross processing artifacts was observed in the SEM micrographs when compared to optical images of the cryptobiotic specimens (**Figure 1**). The same morphological features for each cryptobiotic state present in the optical images were reflected in many of the SEM micrographs obtained, with few added structural alterations that could be amributed to any of the above processes involved in SEM imaging (e.g., limle appearance of apparent collapsed structures from evaporation).

**Figure 1:**
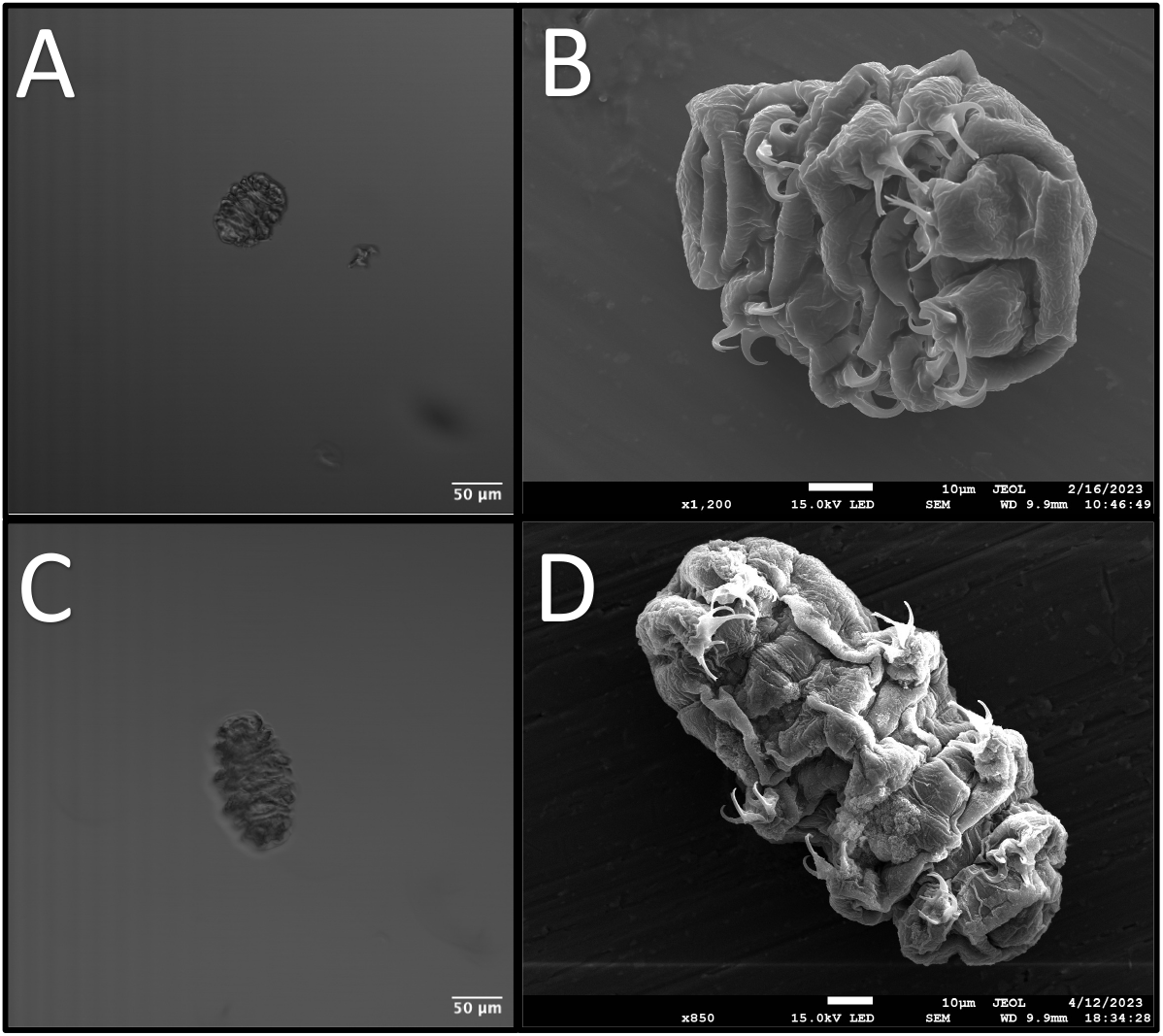
Optical Versus Electron Imaging of Osmobiotes. A – Optical image of a sucrose-induced tun; B – SEM micrograph of a sucrose-induced tun; C – Optical image of a CaCl_2_-induced tun; D – SEM micrograph of a CaCl_2_-induced tun. The overall structure of both kinds of cryptobiotes seems to be preserved through the sample preparation process for SEM analysis).

After the preservation of morphology in sample preparation was confirmed, cryptobiotes formed by both stress conditions were compared to determine the structural differences present between the specimens. The qualitative analysis of the SEM micrographs of sucrose- and CaCl_2_-induced tuns demonstrates a few key distinctions between their overall morphologies (**Figure 2**). First, the diminished contraction along the anterior-posterior axis for CaCl_2_-induced tuns that was observable in optical images (**Figures 1A and 1C**) is confirmed. The micrographs reveal two factors contributing to this: 1) the anterior and poster portions of the CaCl_2_-induced tuns appear to curl towards the main body portion to a lesser extent, and 2) the contraction along the anterior-posterior axis appears to be lower for CaCl_2_-induced tuns, evidenced in part by the decreased depth of folds of the specimens’ flexible cuticles (**Figure 2A-C vs. Figure 2D-F**, feature 2 in blue).

**Figure 2:**
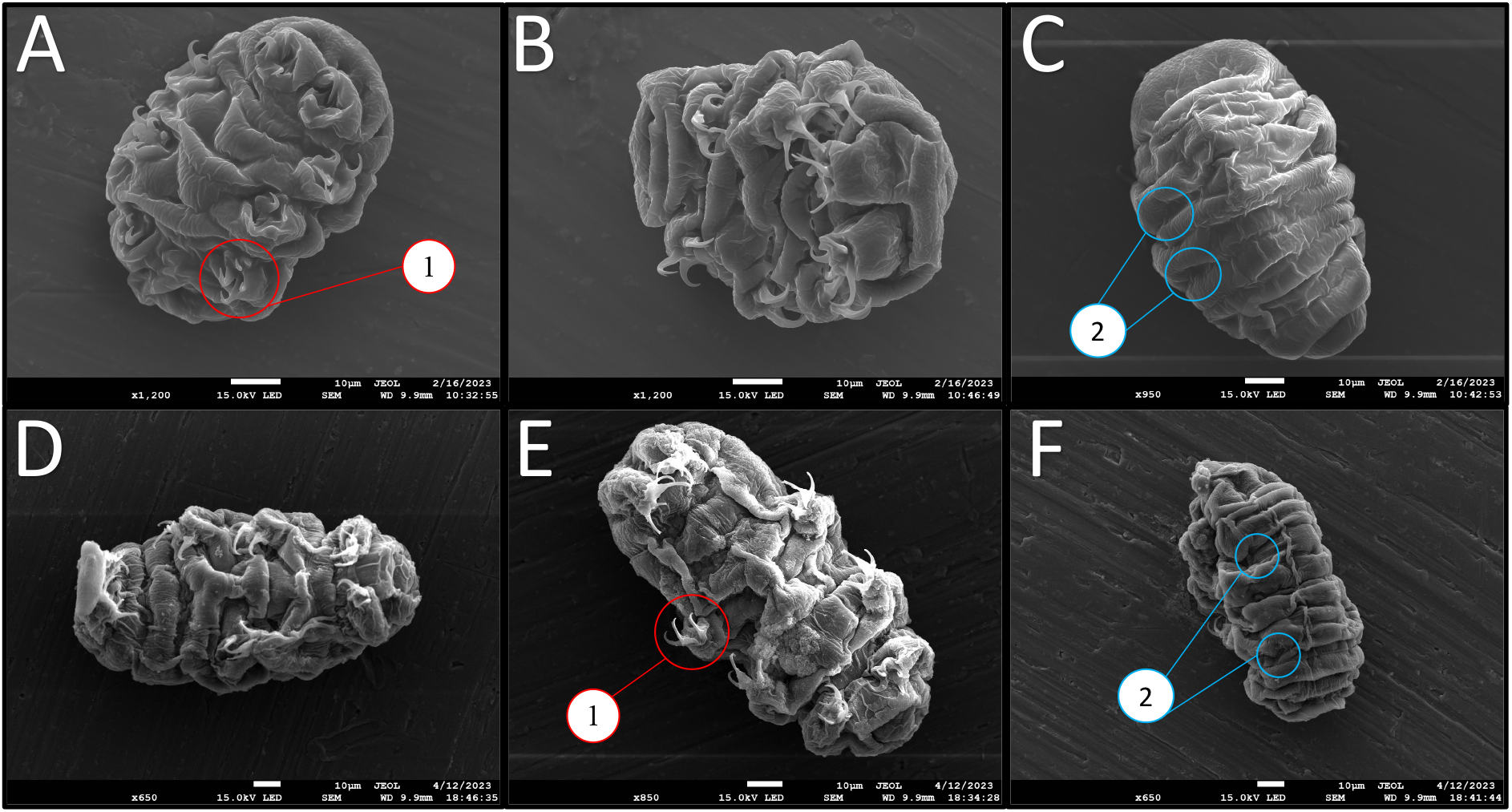
Representative SEM Micrographs Obtained For Sucrose- and CaCl_2_-Induced Tuns. A-C – Sucrose-induced tuns; D-F – CaCl_2_-induced tuns. Panels A/B and D/E show ventral views of the specimens while panels C/F show dorsal views. Two notable features are highlighted by colored/numbered circles. Feature 1 (red) highlights the positioning of the limbs while feature 2 (blue) highlights the differences in the cuticle folds.

Second, the limbs of the CaCl_2_-induced tuns appear to contract into/towards the body cavity less than those of the sucrose-induced tuns. The contraction of the legs all the way into the body cavity observed for the sucrose-induced tun in **Figure 2A** (feature 1 in red) was observed in multiple sucrose-induced tuns, but not observed for any CaCl_2_-induced tun. The limbs of the sucrose-induced tuns also seem to curl towards the midline more than those of the CaCl_2_-induced tuns, whose limbs appear to consistently position along the edge of the body. These distinct differences between the two observed cryptobiotes demonstrate that even very similar stress conditions, such as two distinct tun-inducing osmotic stressors, can induce distinctive morphological alterations that still allow for successful cryptobiosis induction and stress tolerance.

### 3.2. Shadow Imaging by Confocal Scanning Laser Microscopy Shows Volume Differences Between Cryptobiotes

Using a multiphoton (MP) near-infrared laser as the excitation light source along with a cell-impermeant fluorophore for shadow imaging allowed for the reconstruction of full tardigrade specimens, both in their hydrated and cryptobiotic states. Particularly beneficial was the ability of this method to accurately resolve both the limbs and surface contours of the specimens (**Figure 3C**). The axial resolution afforded by the MP laser allowed for the individual acquisition of body components that were stacked on top of one another (**Figured 3D** shows how the limbs were resolved separately from the body cavity). The high concentration of a cell-impermeable fluorophore used generated enough contrast to resolve even the body contours of specimens (**Figure 3E**). The enhanced penetration depth of the MP laser also allowed for resolution of small features (such as limbs) deeper into the sample (**Figure 3F**).

**Figure 3:**
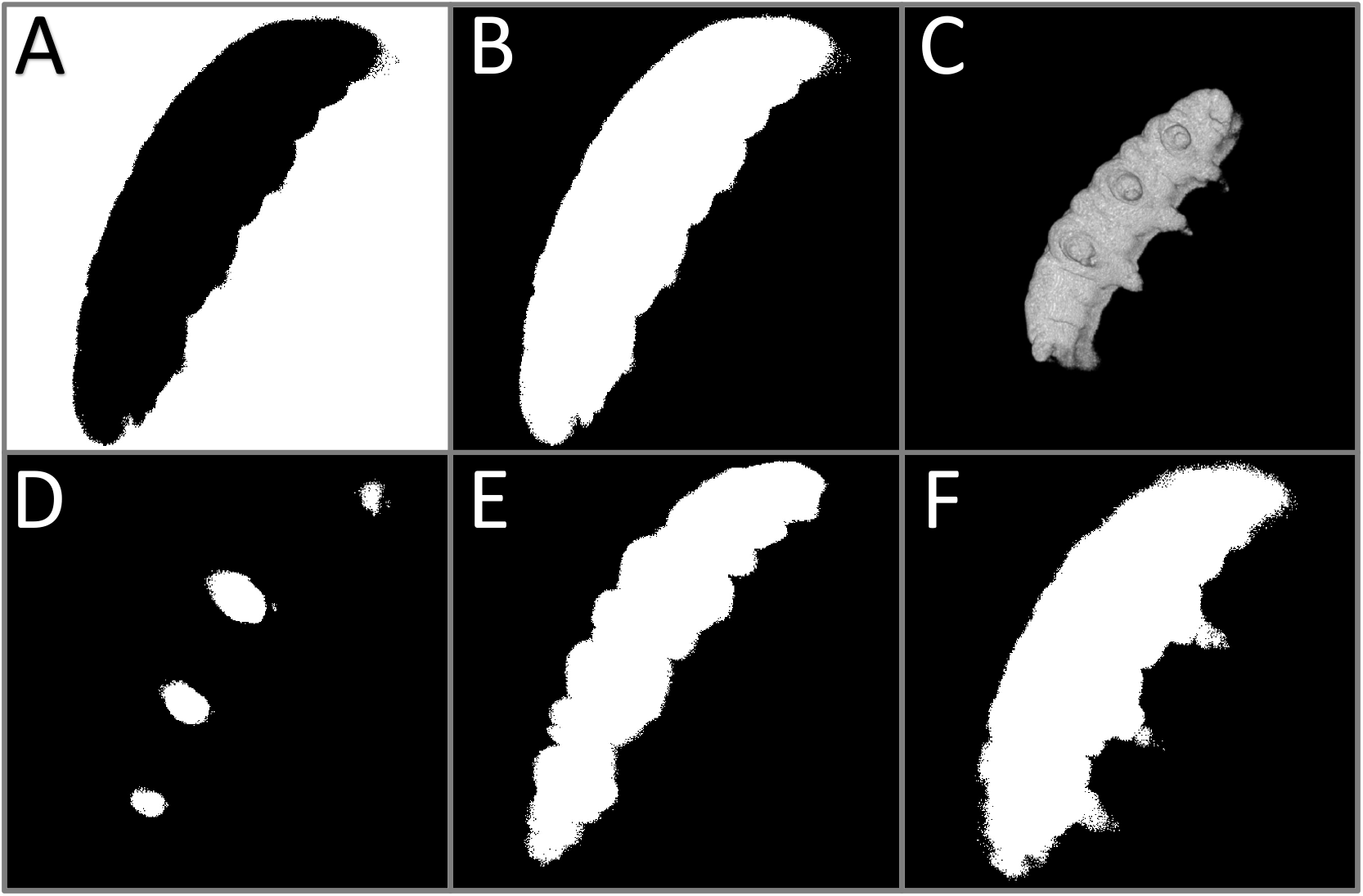
Shadow Imaging Data and Analysis Process. A – Binarized, non-inverted slice; B – Slice after inversion; C – 3D Rendering of inverted stack; D – Slice showing data contributing to the rendering of the limbs furthest from the slide; E – Slice showing data contributing to the rendering of the main body segment, with the cuticular folds evident; F – Slice showing data contributing to the rendering of the body and limbs closest to the slide. Panels A and B and D-F only represent data from individual focal planes. Panels A and B display the same data (same focal plane). Panels D-F are a discontinuous series of slices penetrating deeper into the sample. All data in this figure are from a singular hydrated specimen.

These features of the method allowed for the generation of three-dimensional renderings that possessed nearly all of the features that were observable via SEM imaging, including the cuticular foldings in both the hydrated and tun states (**Figure 4**). Of note for volume estimations is the rendering of the legs of hydrated specimens (**Figure 4A-D**), contributors to overall volume that are often excluded in tardigrade volume estimations. One feature that is noticably absent are the claws that are inserted on the ends of limbs, which likely reflects the fact that the method utilized here cannot resolve sub-micron features such as tardigrade claws (which are sub-micron in diameter). Another point of limitation is that the contact surface between specimens and the slides they are resting on could not be rendered, resulting in flattening of the renderings at that surface (**Figure 4C**). This presumably results in a slight overestimation of volume as the renderings will include the space between the specimen and the slide as part of the specimen. Since the specimens consistently appeared to lay flat against the slides, the added volume likely constitutes the volume of the space between cuticular folds that are touching the slide.

**Figure 4:**
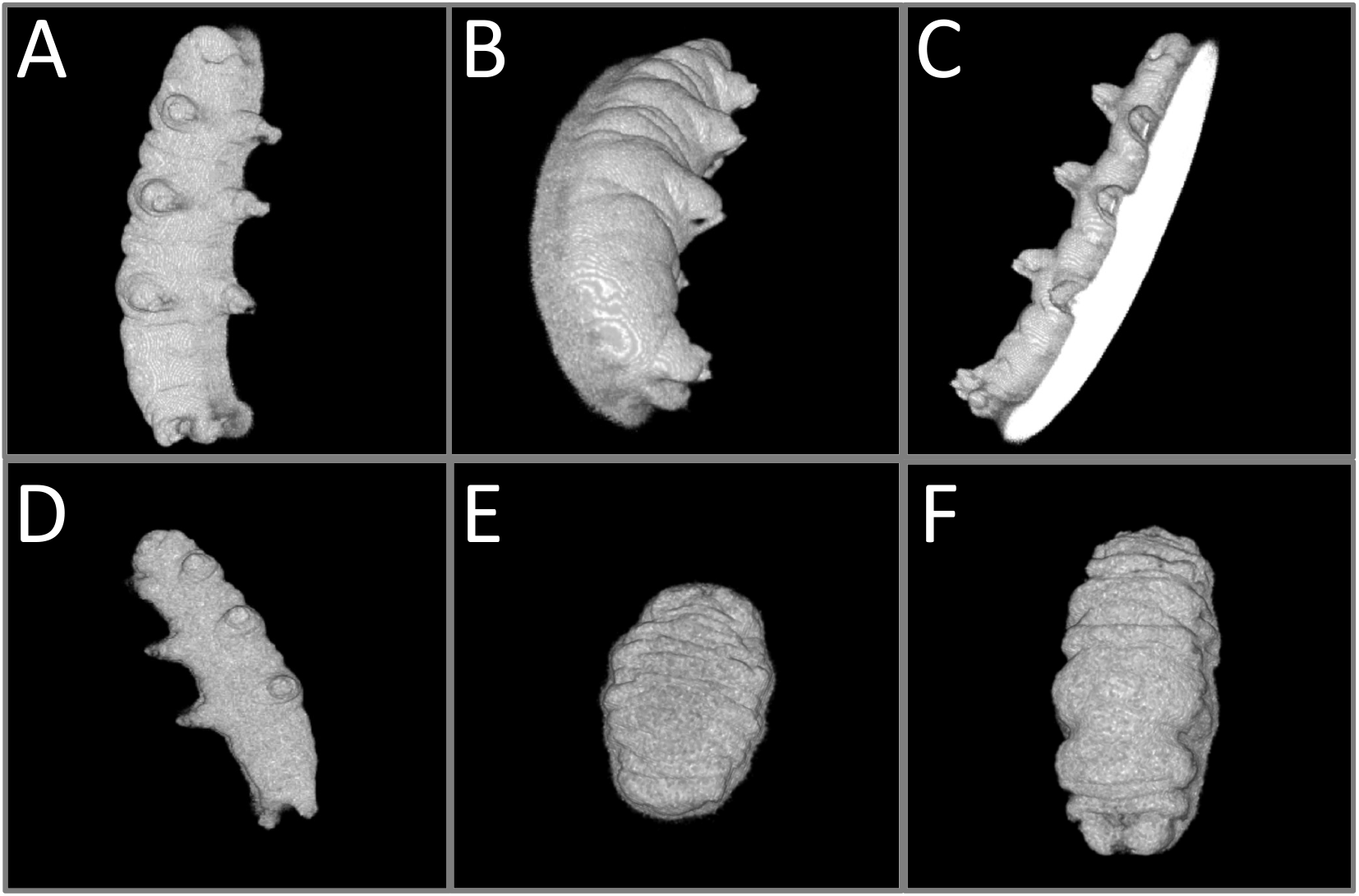
Multiphoton Laser Shadow Imaging Renderings. A – 20° offset from xy view of a hydrated tardigrade; B – 50° offset from xy view of the same hydrated tardigrade; C – View showing the bottom surface of the rendering of the same hydrated tardigrade; D – Rendering of a different hydrated tardigrade; E – Rendering of a sucrose-induced tun; F – Rendering of a CaCl_2_-induced tun.

Overall, the renderings acquired allowed for volume estimations that factored in the contributions of both the limbs and body contours of specimens. These estimations were calculated for hydrated specimens, sucrose-induced tuns, and CaCl_2_-induced tuns. The estimations not only confirmed the significantly reduced volumes of tardigrades in their osmobiotic states in comparison to their normal hydrated state, but also showed statistically significant differences in volume between the two osmobiotic states (**Figure 5**). Hydrated specimens (n = 10) had an average volume of 3.22 × 10^5^ µm^3^ (322 pL), sucrose-induced tuns (n = 10) had an average volume of 1.11 × 10^5^ µm^3^ (111 pL), and CaCl_2_-induced tuns (n = 12) had an average volume of 1.62 × 10^5^ µm^3^ (162 pL). These correspond to sucrose-induced tuns demonstrating a 65.5% reduction in volume and CaCl_2_-induced tuns demonstrating a 49.7% reduction in volume (**Table S1**).

**Figure 5:**
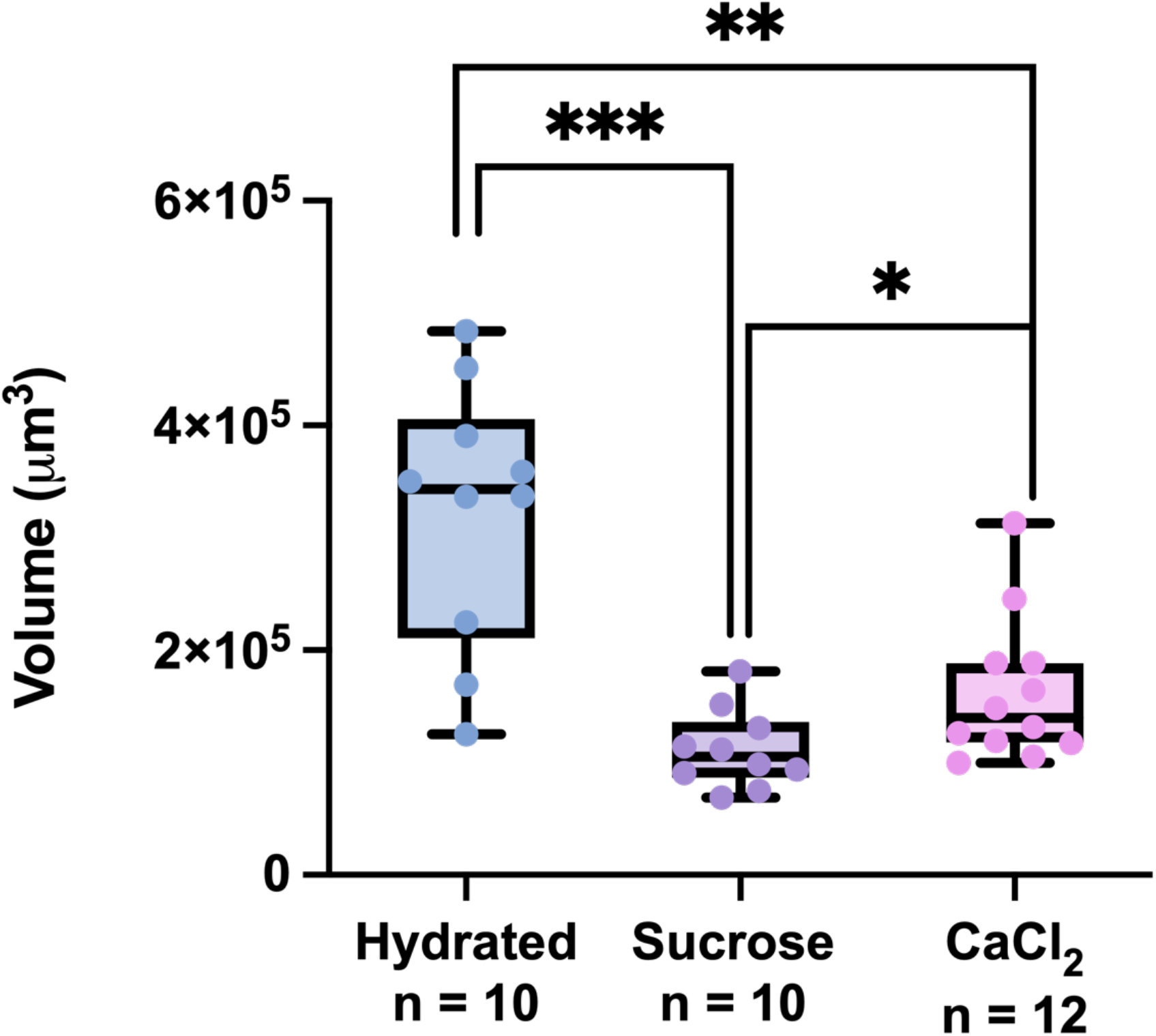
Volume Measurements Made Using Shadow Imaging 3D Renderings. All data points, each of which represents the measurement of one specimen, are shown in addition to the summary statistics represented by the box-and-whisker plots. Statistical significance was determined by Welch’s t-test. * - p < 0.05, ** - p < 0.01, *** - p < 0.0005.

Given that the sucrose (600 mM) and CaCl_2_ (75 mM) solutions used generated different calculated osmotic pressures (14.62 bar and 4.95 bar, respectively), it was initially hypothesized that the measured volume differences were attributable to those differences. To determine if osmotic pressure of the applied osmotic stressors determines/affects resultant tun volumes, samples of sucrose-induced tuns formed in response to two other concentrations of sucrose (300 mM and 450 mM) were measured as well. The average volume of tuns formed in response to 300 mM (П = 7.31 bar, n = 13), 450 mM (П = 10.97 bar, n = 13), and 600 mM (П = 14.62 bar, n = 10) sucrose solutions were 104,000 cubic microns, 135,000 cubic microns, and 111,000 cubic microns, respectively (**Figure 6**). There were no statistically significant differences (p < 0.05) between these groups, and no trend towards increasing volume (less expelled water during tun formation) was observed. Thus, stressor concentration (osmotic pressure) does not seem to substantially influence the volume of tuns formed, pointing towards stressor-dependent mechanisms of tun formation.

**Figure 6:**
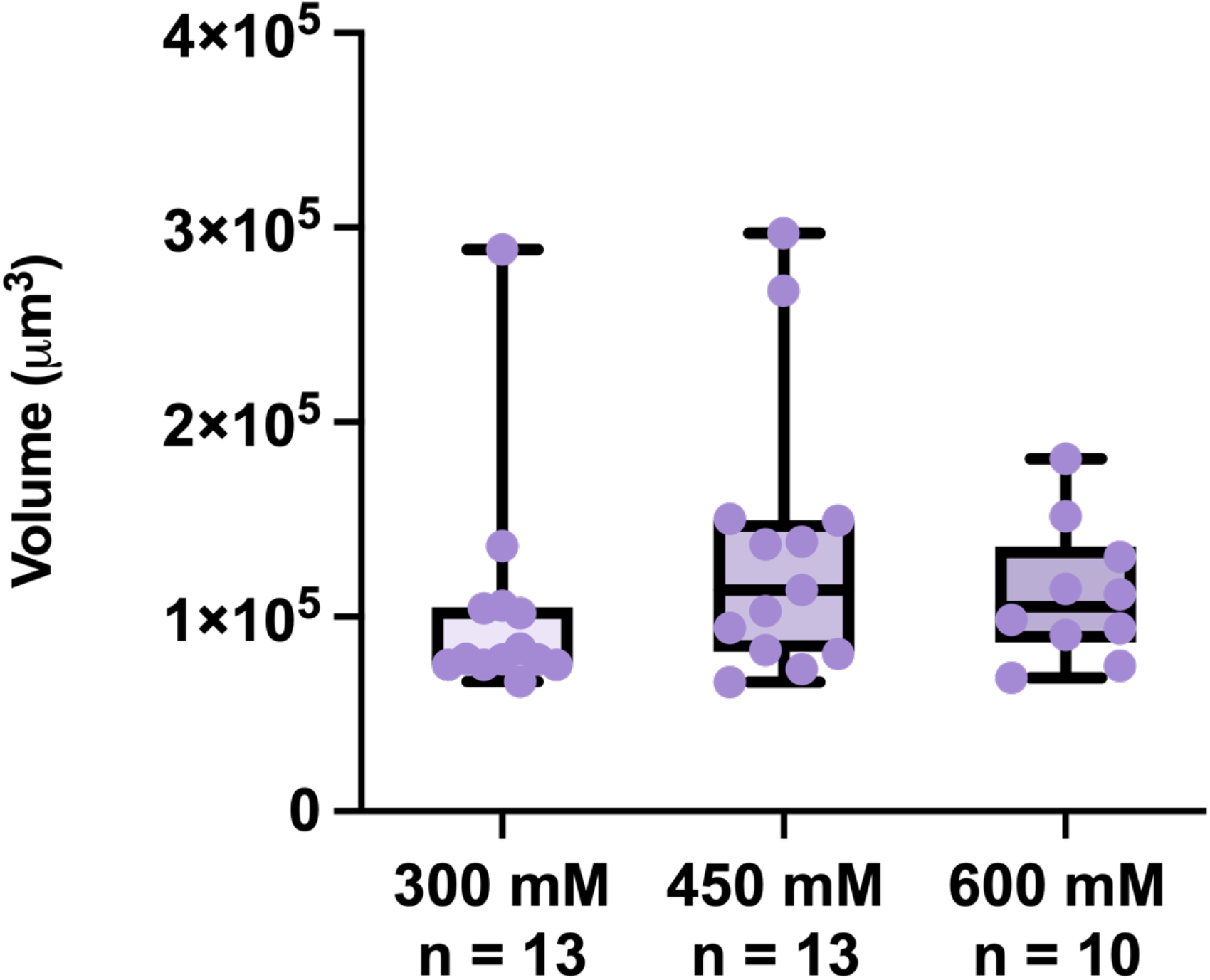
Volumes of Sucrose-Induced Tuns Formed in Response to Varying Concentrations of Sucrose. All data points, each of which represents the measurement of one specimen, are shown in addition to the summary statistics represented by the box-and-whisker plots. Statistical significance was determined by Welch’s t-test. There were no statistically significant differences between groups.

We also imaged a tardigrade that was laying eggs before anesthetization. The rendering produced a relatively good depiction of the event (**Figure 7**), pointing towards a potential for the method described herein to be useful in other contexts.

**Figure 7:**
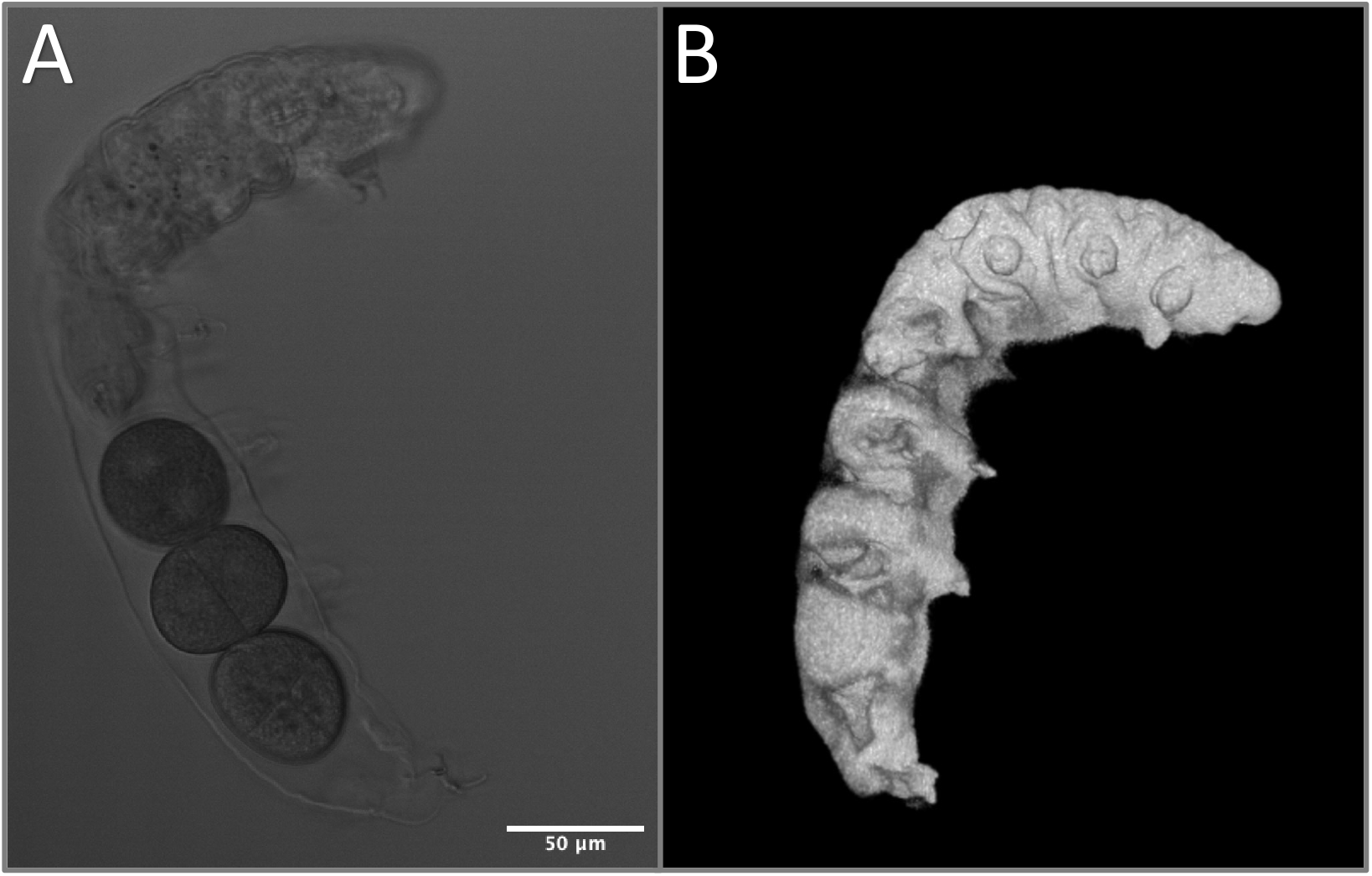
Tardigrade Egg-Laying Visualized by Multiphoton Laser Shadow Imaging. A – One slice of transmitted light stack; B – 3D rendering produced from shadow image stack.

## 4. Discussion

An underexplored area in the investigation of tardigrade cryptobioses is comparing the fine morphological differences between cryptobiotes, especially between cryptobiotes which take on the same general morphology (tun, cyst, turgid state, etc.). The primary limitations for conducting such comparisons are a lack of defined methodologies for comparing fine structural details between cryptobiotes and insufficient methods for quantitatively assessing three-dimensional morphometric data of cryptobiotes, including changes in volume. Elucidating differences between the morphology of cryptobiotes exposed to different stress conditions is essential to understand how tardigrades respond to different stressors, which could reveal different mechanisms of stress sensing and/or tolerance. Herein, a low-cost, accurate, fluorescence-based shadow imaging method was developed and utilized to demonstrate that sucrose- and CaCl_2_-induced osmobiotes are morphologically distinct, providing both a novel method for tardigrade volume estimations as well as insights into how tardigrades respond to osmotic stress. This approach represents a step forward in quantifying morphological differences, opening new avenues of research for researchers to investigate variations in cryptobiosis which may reveal fundamental physiological mechanisms in the induction of cryptobiosis.

SEM was chosen to conduct qualitative analysis due to its abilities to image with high resolution and provide high contrast micrographs of translucent organisms (like tardigrades) which would normally be difficult to characterize with visible light techniques. The sample preparation optimized here represents a simple approach to collecting a reasonable number of high-quality micrographs that can be used downstream for comparing cryptobiotes, opening the door for future investigations into the fine morphological transformations occurring during cryptobiosis induction that may be useful in a variety of research endeavors. However, the methodology presented here will likely not be suitable for all applications. First, we observed structural deformities in hydrated specimens imaged using this “quick” fixation method. **Figure S2A and S2B** show that a one-step fixation and air-drying protocol is not suitable for tardigrades that still possess most of their internal water stores (e.g., hydrated tardigrades). Second, the appearance of similar structural deformities in some of the osmobiotic tuns (**Figure S2C and S2D)** makes the method less suitable for investigations into features that cannot be verified by other techniques as it would be difficult to discern whether any observed differences would be the result of true, controlled morphological transformations or simply a result of artifacts introduced by the sample preparation. Despite these limitations, the method described here did allow for a comparison of wholesale morphological features (the extent of contraction in various body regions) between the two cryptobiotic states investigated. Additional fixatives (glutaraldehyde, osmium tetroxide, and uranyl acetate) as well as different dehydration protocols (critical point drying or the use of hexamethyldisilizane for air-drying) could improve sample preservation during preparation, and others have implemented such approaches for imaging tardigrades via SEM in the past.^11-13, 24^

Despite the aforementioned limitations, SEM micrographs of the osmobiotes facilitated the observation of many morphological differences between the sucrose- and CaCl_2_-induced tuns that would have been difficult to observe with other techniques. First, the diminished contraction of CaCl_2_-induced tuns in comparison to sucrose induced tuns was confirmed, highlighting that differences between the two could be discerned even without the use of higher magnifications. Further, there were apparent differences in muscular coordination between the two sets of osmobiotes. Sucrose-induced tuns displayed significantly more “curling” of their limbs and head/tail regions in forming a more spherical tun while CaCl_2_-induced tuns displayed far less change from the hydrated state (showing little limb retraction and minimal arching of the main body). These differences overall demonstrate that the morphology of the two different sets of osmobiotes are distinct, indicating that there may be differences in how *H. exemplaris* responds to each stressor despite both stressors being putative osmotic stressors. Future investigations into the muscular coordination/activation in both tun states would help to elucidate why the morphological differences appear. The use of calcium-sensitive dyes would be useful for investigating neurological/muscular activity that may differ in intensity or localization between the two conditions. Fluorophore-conjugated phalloidin would allow imaging of the coordination of muscular groups in both states, and that approach has been demonstrated in tardigrades already (though not in *H. exemplaris*).^11^ A general consideration for any future use of these kinds of analyses is that they should be conducted across multiple different stress conditions, including conditions that are generally classified as the same kind of stress (e.g., both sucrose- and CaCl_2_-induced tuns should be analyzed), as is emphasized by the work presented herein.

Beyond qualitative analysis, the work presented here also provides a novel quantitative method for analyses of the three-dimensional features of tardigrades and tardigrade cryptobiotes, particularly their volumes. Building upon previous work describing shadow imaging,^21^ this work developed an easy-to-implement method to produce three-dimensional renderings of tardigrades in both their hydrated and osmobiotic states, allowing for more precise determinations of their volumes than has previously been accomplished in the literature. While the method was initially conducted using an argon laser as an excitation light source, the switch to using a multiphoton laser for fluorophore excitation provided significant retention of fluorescence signal at high sample depths in comparison to the argon laser (**Figure S1**). This switch was motivated by the multiphoton laser’s greater penetration depth and reduced depth of field, thus ensuring less out-of-focus light due to excitation requiring a high photon density.^25^ However, switching to the multiphoton laser does make the method outlined here potentially prohibitive for some investigators as multiphoton (Ti:sapphire) lasers are not as common as gas lasers. This may motivate future work into other alternatives for decreasing signal depletion near the slide. Importantly, we also found that measuring and comparing the volumes of different tardigrade states does not require utilizing the same sample of tardigrades for each measurement; percent differences between states remain consistent whether measured using the same specimens or different ones (**Tables S1 and S2**). This indicates that different samples of tardigrades (at least when using a sample size of ca. 10) tend to have reproducible volumes. This means that measuring the same specimen in different physiological states is not necessary, which is convenient as doing so is far more laborious (each additional specimen analyzed would require another round of plating, measurement, removal from slide, cryptobiosis induction, re-plating, and re-measurement).

Using renderings produced from shadow imaging, volume determinations showed that CaCl_2_-induced tuns have a greater mean volume than sucrose-induced tuns (**Figure 5**), which seems attributable to their comparatively greater length observed via SEM (**Figure 2**). This points towards sucrose-induced tuns having a lower internal water content than CaCl_2_-induced tuns. Further, sucrose-induced tuns formed in response to lower concentrations of sucrose (450 mM and 300 mM) did not have increased volumes in comparison to those formed in response to 600 mM sucrose (**Figure 6)**, indicating that the observed volume differences between sucrose-induced and CaCl_2_-induced tuns were not a result of differences in induced osmotic pressure. Previous work demonstrated that tardigrade water content is likely controlled via active transport rather than passive osmosis.^12^ Therefore, the differences in volume observed here between sucrose- and CaCl_2_-induced tuns—that are not apparently dependent upon the concentration of applied osmotic stressors—indicate that tardigrades respond uniquely to each stressor, suggesting unique stressor-dependent pathways of tardigrade cryptobiosis. This comports with reports that tardigrade survival differs between exposure to ionic and non-ionic osmolytes.^18, 19^ The unique responses to ionic and non-ionic osmolytes raises questions about how tardigrades are responding differently at the cellular level, and future studies should investigate these questions.

Considering that the method for volume measurement presented here may be burdensome or unnecessary for some investigations, we compared the measurements acquired to estimates of the specimens as perfect cylinders/hemicylinders (**Tables S4 and S5**). Those comparisons reveal that the estimations of tardigrades as perfect shapes can result in significant over- and underestimations of volumes depending on the tardigrade state and the method of estimation used (**Table S4**). An important note is that the diameter of tardigrades was not the same in all directions (**Table S3**), and utilizing different axes for the diameter measurement resulted in starkly different volume estimations (**Table S4**). This highlights a substantial limitation of the geometrical approximation approach when three-dimensional data is not acquired: diameter measurements cannot be consistent as the diameter measured will vary depending on the orientation of the specimen (on its side or facing up/down, for instance). This can lead to significant variations in calculated volumes within experimental groups, making it difficult to observe differences between groups. However, calculation of the correlation coefficient, r, between the volumes determined from shadow imaging and the volumes determined through estimations shows that the differences in volume between specimens correlates well between the two methods when the diameter is measured the same way for each specimen. This shows that estimations of volume that have been used previously may be suitable when doing relative rather than absolute quantifications of volume. An important note is that the correlations depend on how the diameters of the specimens are measured (**Table S5**). Correlations are improved when volume estimations measure the diameter as an average of the specimens’ widths in two different dimensions, which is sensible given that the organisms are not perfect cylinders or hemicylinders.

Overall, these analyses indicate that the geometrical approximation method for tardigrade volume estimation does not produce accurate or even necessarily precise results (in comparison to shadow imaging); however, the method can be precise if the diameter is measured in the same way for each specimen (measuring the dorsal-ventral length for each specimen, for instance), making it useful for relative measurements such as percent differences (though this requires either only measuring specimens that are oriented the same way or collecting three-dimensional image data).

One limitation of the shadow imaging method presented here is that it can’t produce complete three-dimensional renderings of specimens. At the contact surface between the specimen and the slide, we observed fluorescent signal depletion, causing the renderings to show flattening at that contact region (**Figure 4C**). This adds to measurement error when the renderings are used to calculate the volumes of imaged specimens. However, this error shouldn’t be prohibitive of the use of the shadow imaging method for the following reasons: 1) The error in the contact surface rendering should be relatively consistent across all datasets, preventing the artifact from skewing the data, 2) The contact surface should only represent a very small volume fraction, meaning the error added to measurements should be minimal, and 3) Such an artifact is present in other microscopic methods for volume measurements, such as in volume electron microscopy.^26^ Overall, the method described is useful both in qualitative (as morphology is accurately rendered) and quantitative (volume determinations) analyses.

Finally, the ability of the multiphoton-based shadow imaging to produce an accurate rendering of tardigrade egg-laying indicates that this method is not restricted solely to measuring the volumes of tardigrades (**Figure 7**). Other specimens may very well be suitable to analyze via this method. The general consideration to be made in determining suitability is the permeability of the exterior structure of the specimen of interest to the fluorophore being used. A cell-impermeant fluorophore (calcein) coupled with the low permeability of the tardigrade cuticle to bulky molecules made this method excellent for the work described here, but additional considerations may be necessary in other applications of the method.

Overall, the work presented here demonstrates that tardigrade cryptobiotes induced by different osmotic stressors can form unique tuns, the volumes of which are not apparently dependent upon the concentration of the stressor used. This indicates that tardigrades may actively regulate their internal water content and demonstrates that they can respond differently to distinct osmotic stressors. This work also provides steps forward in pursuing more rigorous qualitative and quantitative analysis of tardigrade cryptobioses which will be essential in pursuing questions related to the mechanisms of tardigrade stress tolerance and stress sensing. Further, the work conducted on developing multiphoton-based shadow imaging may prove useful in other fields of research.

## Acknowledgements

The authors thank the Marshall University Molecular and Biological Imaging Center for the use of its microscopy equipment for this investigation.

## Competing Interests

The authors declare no conflicts of interest.

## Author Contributions

Conceptualization, A.S., B.F., D.K., H.O., K.J. and L.H.; methodology, B.F., D.N., and M.N.; validation, B.F.; formal analysis, B.F.; investigation, B.F., H.O., and K.J.; resources, D.K., L.H., and M.N.; data curation, B.F.; writing—original draft preparation, B.F.; writing—review and editing, A.S., B.F., D.K., D.N., L.H. and M.N.; visualization, A.S., B.F., and K.J.; supervision, D.K., D.N., L.H., and M.N.; project administration, D.K., D.N., L.H., and M.N.; funding acquisition, D.K., L.H., and M.N. All authors have read and agreed to the published version of the manuscript. Author contributions are listed in alphabetical order by first initial and then by last initial where necessary.

## Funding

This research was supported by National Science Foundation grants awarded to L.M.H. (NSF-MCB 2149172) and D.R.J.K. (NSF-MCB 2149173). A.L.S. acknowledges funding from the North Carolina Space Grant. Marshall University students were supported by the NASA West Virginia Space Grant Consortium (Grant no. NNX15AK74A). The Leica SP5 confocal and JEOL JSM 7200 FLV microscopes were obtained under National Science Foundation grant award numbers (0959012) and (1828358), respectively.

## Data Availability

Raw numerical data has been provided in the supplementary information for this manuscript.

## Notes

### Competing Interest Statement

The authors have declared no competing interest.

